# Substantial Batch Effects in TCGA Exome Sequences Undermine Pan-Cancer Analysis of Germline Variants

**DOI:** 10.1101/445049

**Authors:** Roni Rasnic, Nadav Brandes, Or Zuk, Michal Linial

## Abstract

**Background:** In recent years, research on cancer predisposition germline variants has emerged as a prominent field. The identity of somatic mutations is based on a reliable mapping of the patient germline variants. In addition, the statistics of germline variants frequencies in healthy individuals and cancer patients is the basis for seeking candidates for cancer predisposition genes. The Cancer Genome Atlas (TCGA) is one of the main sources of such data, providing a diverse collection of molecular data including deep sequencing for more than 30 types of cancer from >10,000 patients.

**Methods:** Our hypothesis in this study is that whole exome sequences from healthy blood samples of cancer patients are not expected to show systematic differences among cancer types. To test this hypothesis, we analyzed common and rare germline variants across six cancer types, covering 2,241 samples from TCGA. In our analysis we accounted for inherent variables in the data including the different variant calling protocols, sequencing platforms, and ethnicity.

**Results:** We report on substantial batch effects in germline variants associated with cancer types. We attribute the effect to the specific sequencing centers that produced the data. Specifically, we measured 30% variability in the number of reported germline variants per sample across sequencing centers. The batch effect is further expressed in nucleotide composition and variant frequencies. Importantly, the batch effect causes substantial differences in germline variant distribution patterns across numerous genes, including prominent cancer predisposition genes such as BRCA1, RET, MAX, and KRAS. For most of known cancer predisposition genes, we found a distinct batch-dependent difference in germline variants.

**Conclusion:** TCGA germline data is exposed to strong batch effects with substantial variabilities among TCGA sequencing centers. We claim that those batch effects are consequential for numerous TCGA pan-cancer studies. In particular, these effects may compromise the reliability and the potency to detect new cancer predisposition genes. Furthermore, interpretation of pan-cancer analyses should be revisited in view of the source of the genomic data after accounting for the reported batch effects.

## INTRODUCTION

Identifying predisposition variants underlying cancer heritability is of utmost importance and a critical milestone for personalized medicine. Despite the enormous clinical relevance, strong evidence for variant contribution to cancer development is restricted to only a few genes harboring significant effects. For example, inherited mutations in BRCA1 and BRCA2 predispose to very high risks for breast and ovarian cancers [1-3]. The risk and prevalence of specific germline variants in cancer predisposition genes greatly vary across ethnicities and cancer types, as illustrated by the high prevalence of BRCA1 variants in Ashkenazi Jews [4, 5]. While each cancer type may have its own signature, a substantial overlap in the identity of known predisposition genes has been observed [6, 7]. Studies of families with high recurrence of cancer identified numerous genes carrying germline mutations with high penetrance (e.g., [2, 8]). The increasing number of sequenced exomes has led to the discovery of additional cancer predisposition genes, mostly with rare mutations [9-11].

In recent years, the task of identifying predisposition variants [2] using data-driven and statistically-sound approaches has become feasible, thanks to the availability of thousands of genomic samples with satisfying sequencing depth and quality, from healthy and diseased individuals (e.g., [7, 12]). The premise is that identifying germline cancer predisposition genes will lead to improved clinical diagnosis of hereditary cancers [13]. The Cancer Genome Atlas (TCGA) [14] is the most exhaustive collection of such data. Batch effects in miRNAs-Seq, RNA-Seq and DNA methylation data from TCGA were reported [15]. However, batch effects in genomic data from whole exome sequencing (WES) were mainly attributed to platform-dependent sequencing reactions and sampling conditions [16]. Additionally, it was noted that TCGA exome sequencing data is liable to inaccuracies resulting from sample calling quality [17] and additional technical effects associated with different batches [18]. The latter is evident through monitoring loss of function (LoF) mutations, and specifically short indels that cause frameshifts [18].

In this study, we performed a detailed analysis of germline variants (common and rare) across six cancer types covering thousands of samples. Our assumption is that germline variants identified using WES of healthy blood samples extracted from cancer patients are not expected to show systematic differences across cancer types, assuming that biases attributed to variant calling, indel recording, and population structure are eliminated. Consequently, the reliability and consistency of the data in TCGA can be directly assessed in an analysis avoiding or correcting for such known confounders. In this study, we show that the mapped reads are already subjected to substantial batch effects, and demonstrate the impact of such batch effects on critical statistical measures of the data and pan-cancer downstream interpretation.

## RESULTS

### Germline variants in exome sequences

In order to test the TCGA dataset for potential batch effects, we processed and analyzed a subset of the cancer-type cohorts in TCGA. We focused on six cancer types, each with at least 250 germline samples (total of 2,241 samples): BRCA (Breast Invasive Carcinoma), UCEC (Uterine Corpus Endometrial Carcinoma), STAD (Stomach Adenocarcinoma), SKCM (Skin Cutaneous Melanoma), LIHC (Liver Hepatocellular Carcinoma) and THCA (Thyroid Carcinoma) (Supplementary Table S1).

We implemented a unified variant calling pipeline for aligned reads (i.e., TCGA germline BAM files) using conventional, well-accepted variant calling methods (see Methods). We restricted the reported analysis to 1,522 samples TCGA classified as Caucasian (marked “White” by TCGA) to eliminate possible biases due to ancestry. We also restricted our analysis to samples profiled using Massively Parallel Sequencing (MPS) methodology (only HiSeq) to minimize variations due to the technical genomic data production protocols. As short indels account for the majority of batch effects and inconsistencies [18], they were not included in the variant calling, and only Single Nucleotide Variants (SNVs) were considered.

### Batch effects manifestation in the number of called variants

Our quantitative analysis reveals a significant batch effect in the number of germline variants per sample across different cancer types. The most prominent characteristic shared by cancer types with similar numbers of called variants is the sequencing center contributing to the collection in TCGA (Figure 1A). The healthy blood samples from patients with skin, stomach and thyroid cancers (SKCM, THCA and STAD) were sequenced at the Broad Institute (BI); samples from patients with uterus and breast cancers (UCEC and BRCA) were sequenced at the Washington University Genome Sequencing Center (WUGSC) and samples from lung cancer patients (LIHC) were sequenced at the Baylor College of Medicine (BCM) sequencing center.

**Figure 1.**
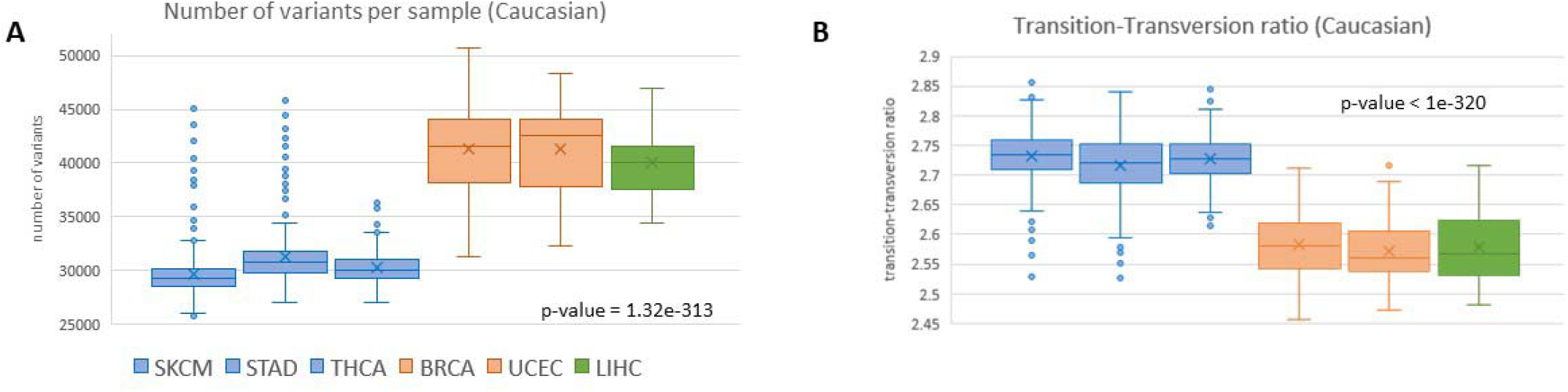
Variability in called variants across TCGA sequencing centers. Batch effect due to sequencing center in 1,522 samples associated with Caucasian populations (originated in Europe, Middle East or North Africa) across the six analyzed cancer types. **(A)** Number of called exome variants per sample. **(B)** Ratio of transition-transversion (TITv) variants per sample. Colors represent the genomic sequencing centers: BI (blue), WUGSC (orange) and BCM (green).

Numerous aspects of the data analysis are sensitive to the origin of the data, thus reflecting the effect of the different batches. We present several such quantitative measures:

### 20-30% difference in the number of called variants per sample

In Figure 1 we show the number of called variants per sample, partitioned by the patient’s cancer type. The average number of germline variants greatly varies across sequencing centers. Samples provided by WUGSC and BCM have up to 30% more variants compared to samples provided by BI (one-way ANOVA, p-value = 1.32E-313). This observation applies also to the other ethnic groups (Supplementary Figure S1).

Recently, a report on a catalogue of rare pathogenic germline mutations from TCGA was presented [7]. This report relied on a different variant calling pipeline. By extracting the number of variants per sample from this report, we show that our reported batch effect is insensitive to the underlying variant calling pipeline. Supplementary Table S2 provides estimated values for the average number of variants per sample across all 33 cancer types in TCGA extracted from this report. In addition to the three sequencing centers covered in this work, the extracted data also includes a fourth sequencing center, the Sanger center. The overlooked dominating signal of the identity of the sequencing center applies in the data extracted from this report, and generalizes to all 33 cancer types (Supplementary Figure S2A). For the six shared cancer types, we report an almost perfect correlation (r=0.91) between the average number of variants per sample calculated in our analysis to these numbers extracted from the report (Supplementary Figure S2B). We conclude that the reported sequencing batch effect dominates the results regardless of the variant calling pipelines used.

### Variations in nucleotide substitution ratios

We find strong evidence for batch effect in the transition-transversion (TiTv) ratios of called variants per sample across sequencing centers (Figure 1B, one-way ANOVA p-value < 1E-320). Samples sequenced at BI have ^~^6% higher transition-transversion ratio (average 2.73) compared to samples from the other two sequencing centers (average 2.57).

### Variant density per gene

We addressed the possibility that the found differences in the number of variants according to the different batches (Figure 1A) might reflect a naive scaling issue due to different sequencing depths or significance thresholds in variant calling among the sequencing centers. We tested whether the batch effects apply also to the relative variant densities across genes. For each combination of gene and sample, we calculated variant density per nucleotide (i.e., the number of variants divided by the full transcript length). We then calculated Pearson’s correlation for each pair of samples across all transcripts. We show the resulted correlations for the 1,522 Caucasian samples (Figure 2). We find that the batch effect dominates these variant densities, with a strong similarity among samples sequenced at the BI. The lung cancer (LIHC) samples, which were sequenced at BCM, show the largest deviation. We conclude that there are consistent variations among samples from different sequencing centers that are more substantial than naive scaling, leading to enrichment or depletion of called variants in specific genes.

**Figure 2.**
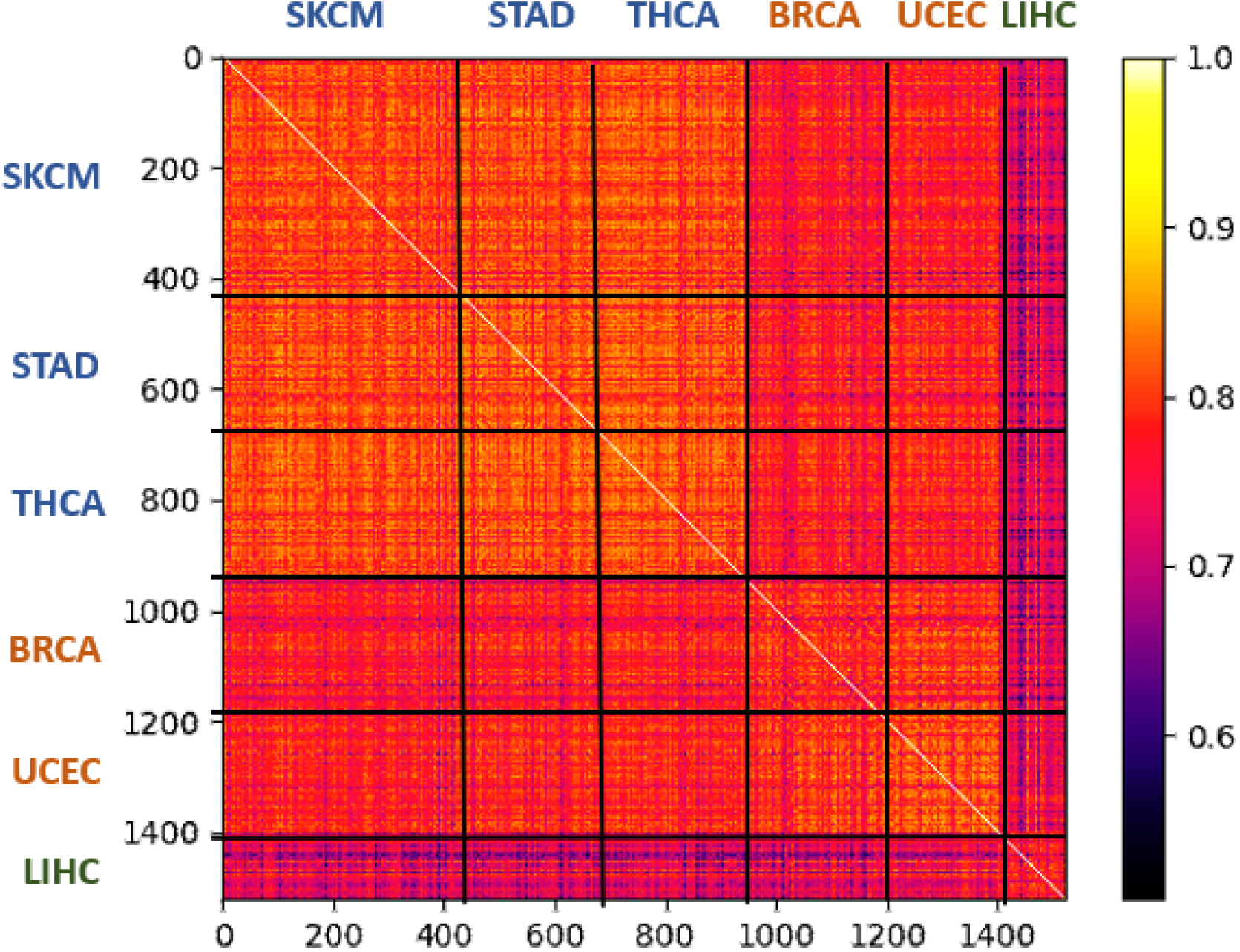
Correlations in variant densities among samples from Caucasian populations. Heatmap of Pearson’s correlations of per-gene variant densities between pairs of samples from Caucasian populations. The 1,522 samples are sorted by their cancer types. The correlation values show high similarity among cancer types sequenced by the same canter. The cancer types are as reported in Figure 1. BRCA (Breast Invasive Carcinoma, 291 samples), UCEC (Uterine Corpus Endometrial Carcinoma, 169 samples), STAD (Stomach Adenocarcinoma, 248 samples), SKCM (Skin Cutaneous Melanoma, 435 samples), LIHC (Liver Hepatocellular Carcinoma, 146 samples) and THCA (Thyroid Carcinoma, 258 samples). Color for the sequencing centers are as in Figure 1.

### Variant distribution within cancer predisposition genes

We tested whether sequencing center batches affect not only the number of variants (Figure 1) and their distribution *among genes* (Figure 2), but also their distribution *within* genes. Figure 3 shows the positional distribution of variants within four known cancer predisposition genes: BRCA1, BRCA2, KRAS and RET. We show marked differences associated to the sequencing centers in the distribution of variants along three of these genes. Interestingly, the strongly reported predisposition gene BRCA2 is mostly indistinguishable for all six cancer types and is thus relatively insensitive to this batch effect.

**Figure 3.**
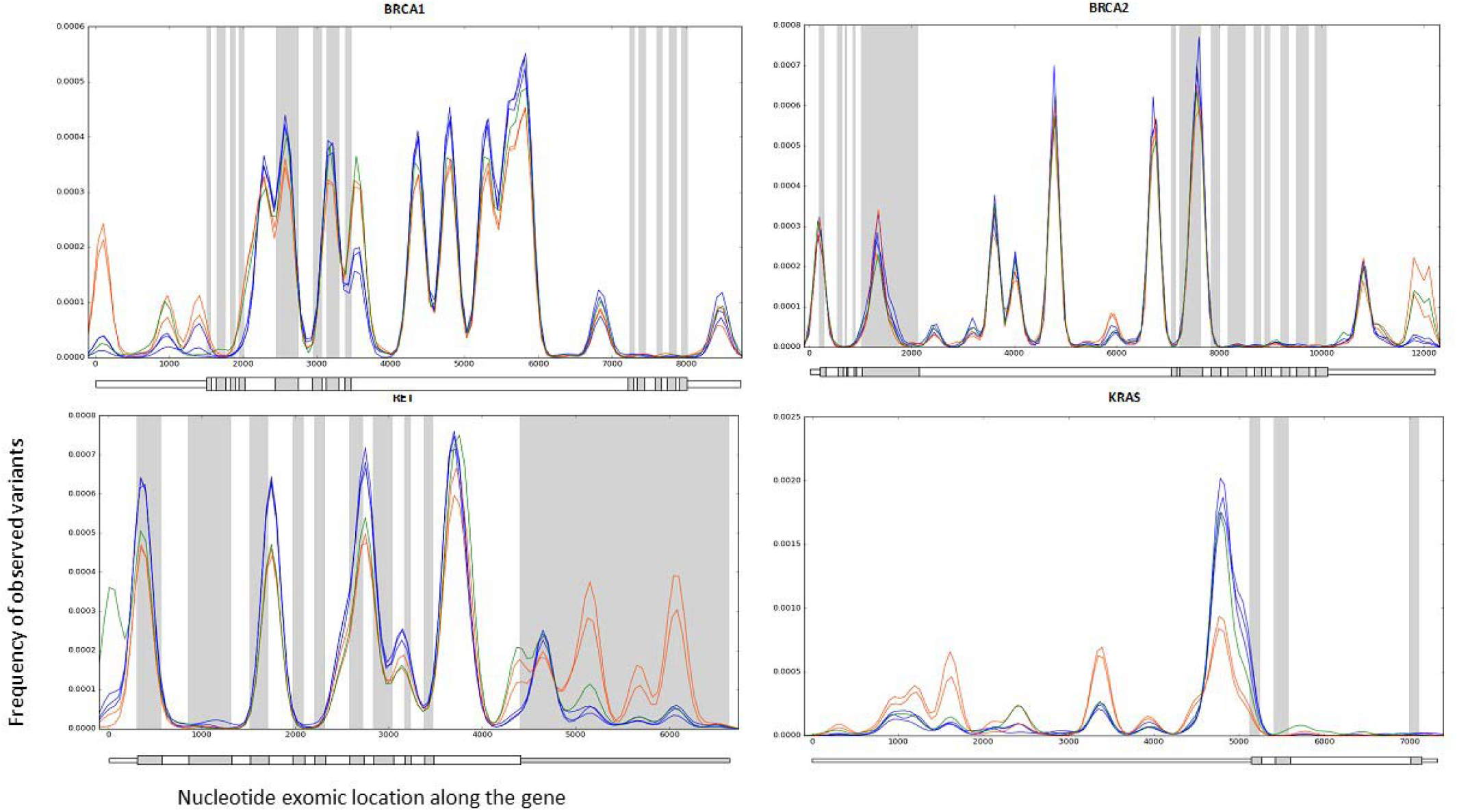
Gene exomic location distributions of germline variants within selected cancer predisposition genes. Empirical probability density functions (PDF) of germline variant coordinates, plotted for four selected genes (BRCA1, BRCA2, RET and KRAS). Each line represents the density function of one of the six cancer types, colored by their corresponding sequencing center: BI (SKCM, STAD and THCA) in blue, WUGSC (BRCA and UCEC) in orange, and BCM (LIHC) in green. The genes BRCA1, RET and KRAS display distinct distributions per sequencing batch, while BRCA2 displays a relatively cohesive distribution. Exons are colored by alternating gray and white backgrounds to enhance the visibility of exon boundaries (introns, for which we have no data, are omitted). The schemas of the transcripts (including the non-coding 5’-UTR and 3’-UTR parts) are shown below each figure. For visibility, the graphs are smoothed by kernel density estimation (KDE), using a window size of 100 nt.

As illustrated in Figure 3 for four selected genes, we analyzed the entire collection of 104 known cancer predisposition genes from the COSMIC catalogue [19], in order to thoroughly quantify the batch effect on the distribution of called variants within those genes (Figure 4). We used the Kolmogorov-Smirnov (KS) statistical test to compare these distributions between the 15 pairs of the six cancer types for each gene. We clustered the genes and pairs of cancer types based on these statistical results (p-values) using Bi-clustering. Pairs originating from the same sequencing center were clustered together (e.g., the three leftmost columns corresponding to the three pairs sequenced at BI), highlighting the effect of the sequencing center on variants’ positional distribution.

**Figure 4.**
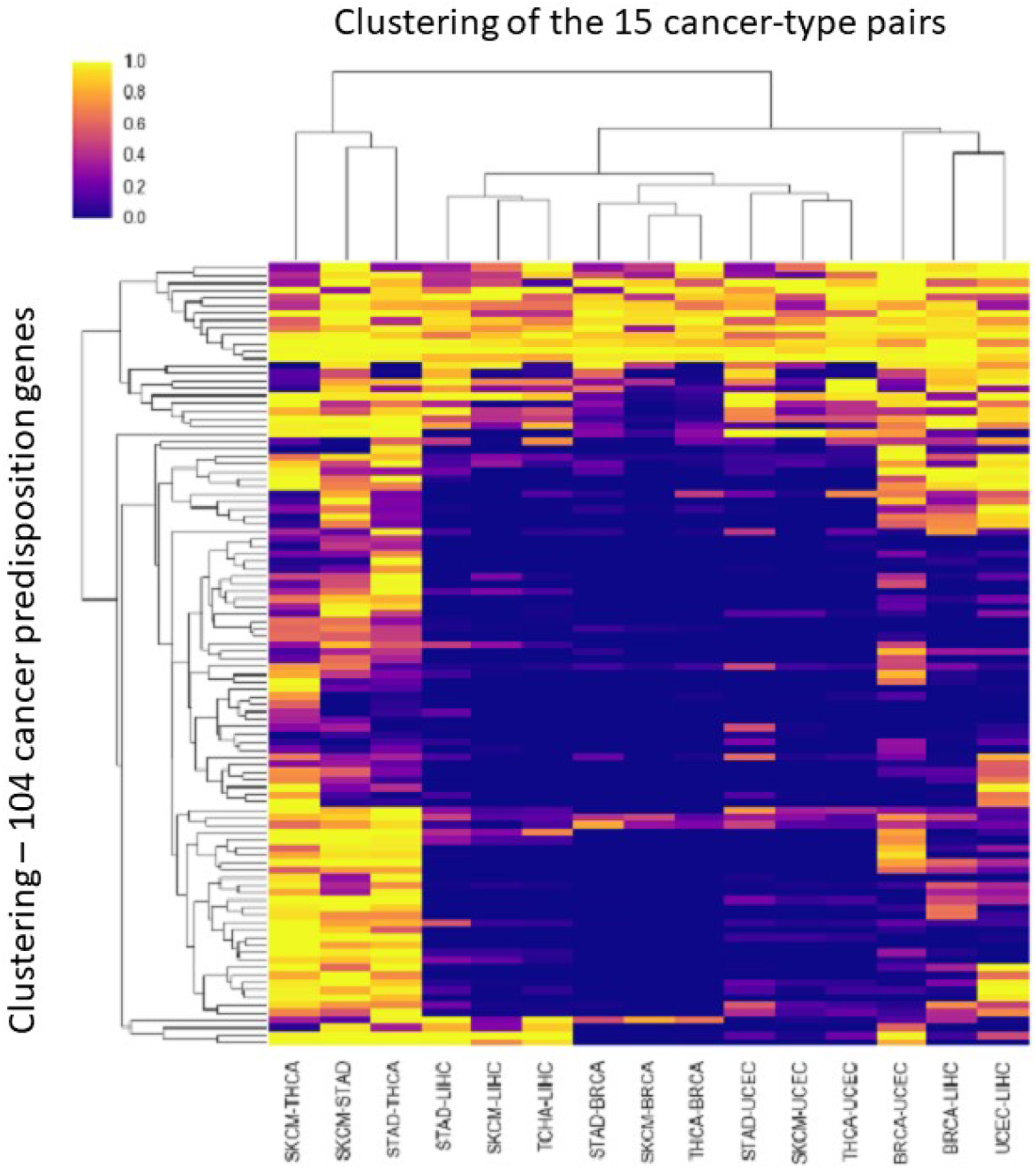
Significant differences in variant location distributions between cancer-type pairs. Bi-clustering of Two-sided Kolmogorov-Smirnov (KS) test results (log-p-values) comparing cancer-type pairs across 104 genes annotated by COSMIC as cancer predisposition genes.

The susceptibility of genes to the batch effect was determined by the ratio of similarity (using KS p-values) *within and across* the BI and WUGSC batches (see Methods). Only 35 of the 104 genes were unaffected by the batch effect (p-value ratio <1.0; Supplementary Table S3). Variant distribution of genes that are extremely sensitive to the batch effect based on this p-value ratio (e.g. MAX, SMACRE1) and other genes that are insensitive (e.g. POLD1) is shown in Supplementary Figure S3.

### Batch effects is associated with clinical outcome

We assume that if the identity and distribution of called variants along genes have no impact on pan-cancer downstream clinical interpretation, there will be no difference between genes that are prone to such batch effect and those that are unaffected by it. To test this assumption we performed an indirect test and followed the survival of patients while focusing on two disjoints gene sets from the 104 genes annotated by COSMIC [19] as germline-associated cancer predisposition genes. Specifically, we sorted the 104 genes by their p-value ratio and defined two extreme gene sets: (i) the top 10% (10 genes) that display maximal sensitivity to the batch effect according to the p-value ratio: MAX, RET, ERBB4, TSC1, DICER1, BARD1, ERCC5, PRKAR1A, PHOX2B and SMARCE1 and (ii) the bottom 10% (10 genes) showing the minimal sensitivity to such effect: CYLD, POLD1, SMAD4, TSHR, CDC73, NTHL1, SMARCB1, TSC2, FH and SDHD. We performed a survival analysis on cancer patients with somatic mutations from an independent cohort, taken from MSK-IMPACT clinical sequenced samples (MSKCC, [20]), which covers 10,129 samples.

We found a clear difference in the Kaplan-Meier estimated survival curves for the two sets of genes (compare Figures 5A to 5B). Specifically, statistically significant reduced survival (Log rank test p-value = 4.58e-4, Figure 5A) is associated with patients carrying mutations in the genes that are maximally sensitive to the batch effect. Such difference is not detected for genes that are resistant to the batch effect (p-value = 0.236, Figure 5B). In both instances, the fraction of cases with mutations in the gene sets is 11% of all 10,129 samples, showing that the difference in observed effects on survival for the two groups of genes is not due to differences in statistical power. We conclude that the relative sensitivity of genes to batch effect may be carried on to downstream analysis, including clinical outcome and its interpretation, even when one uses independent cohorts for such analysis.

**Figure 5.**
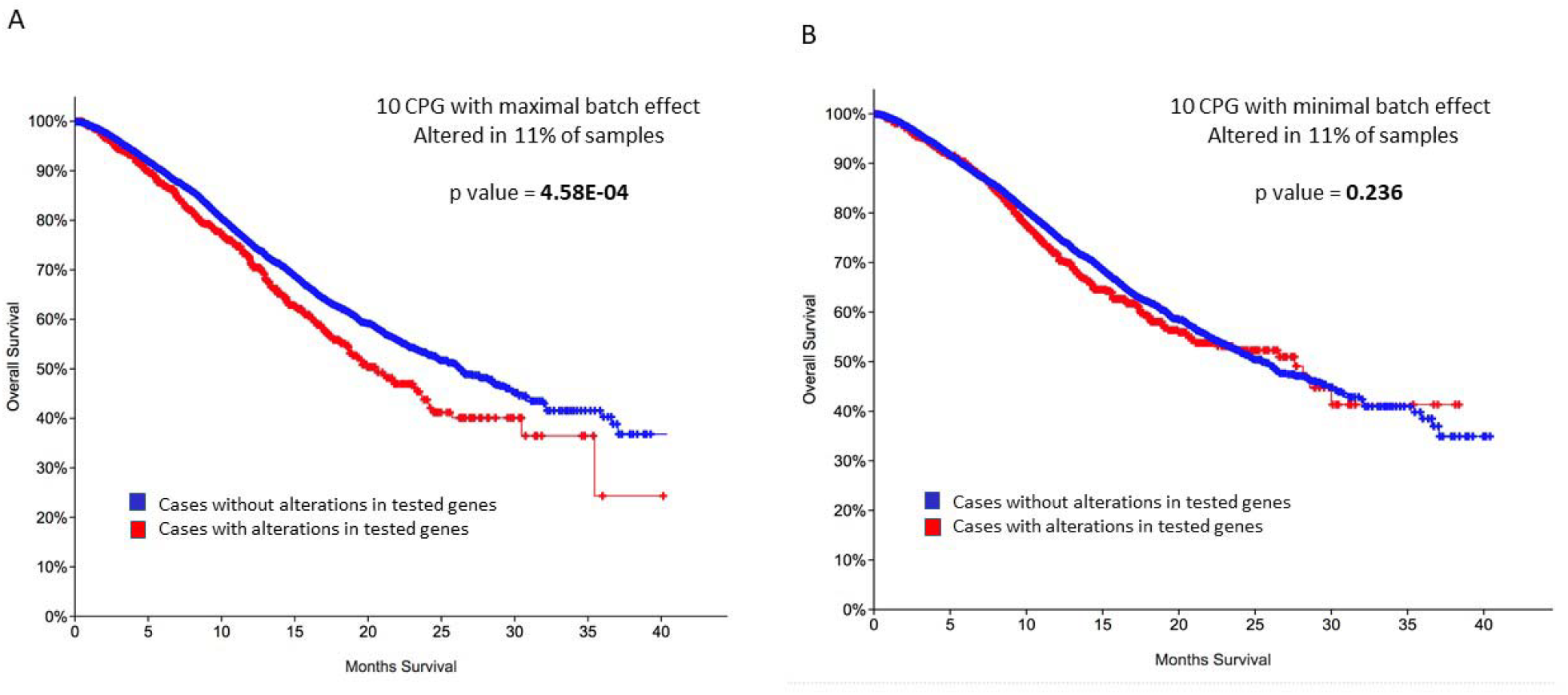
Survival curves for gene sets differing by their sensitivity to batch effect. Kaplan-Meier estimate survival curves tested on 10,129 samples from the MSK-IMPACT clinical sequenced cohort (MSKCC, [19]). The analysis applies to genes from a collection of 104 CPGs annotated by COSMIC. **(A)** Top 10 genes exhibiting maximal sensitivity to the batch effect. **(B)** Bottom 10 genes exhibiting minimal sensitivity to the batch effect. Supplementary Table S3_104 CPG lists the 104 genes along with their batch effect measure.

### Batch effects are associated with most of the analyzed genes

We expanded the KS paired statistics analysis to include all genes with variants in all six cancer types (overall, 18,421 genes). Only 33% of the genes appear to be insensitive to the batch effect (score <1; see Methods and Supplementary Table S3, all genes). Again, we observe strong similarity between cancer-type pairs sequenced at the same centers, compared to high variability between pairs originating from different sequencing centers. Pairs comparing cancer types from BCM (LIHC) and WUGSC (UCEC and BRCA), as well as the UCEC-BRCA pair show intermediate resemblance (Figure 6).

**Figure 6.**
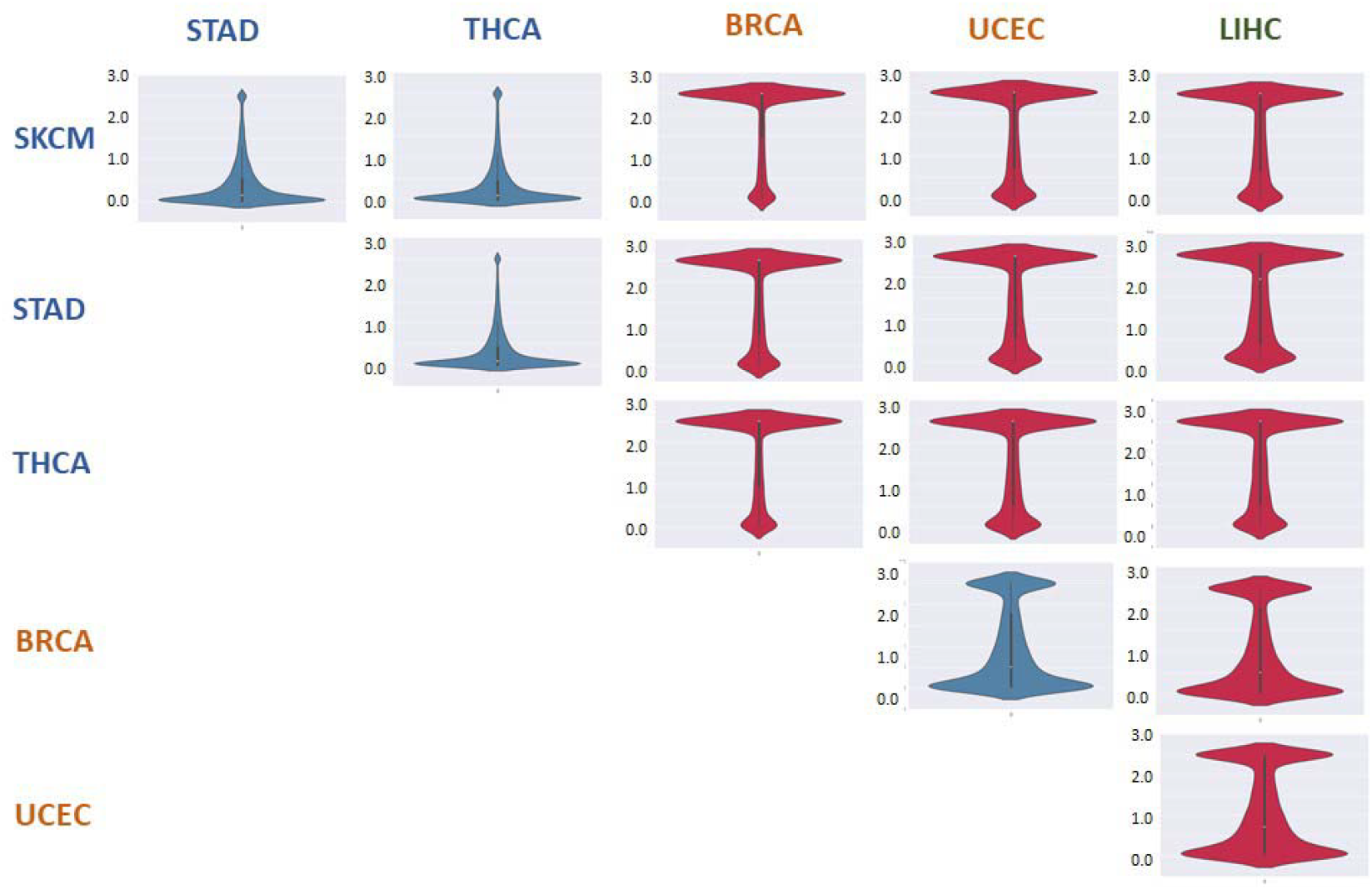
Violin plots based on Kolmogorov-Smirnov test per each group pairing. Two-sided Kolmogorov-Smirnov (KS) tests were carried per gene to test for differences in the distributions of variants between each of the 15 cancer-type pairs (the same variant distributions shown in Figure 3 for four selected genes). Each panel displays the distribution of resulted p-values across all 18,421 analyzed genes. Red-colored images represent cancer-type pairs originating from different sequencing centers, while blue-colored images represent pairs originating from the same sequencing center. Cancer-type labels are color-coded by sequencing centers, as in all previous figures. The y axis scale is -log10(p-value), where all values above 2.5 were truncated to 2.5 (for visibility).

## Discussion

We report on multiple layers of batch effects associated with the sequencing centers contributing to TCGA, which are evident upon examination of called germline variants from thousands of samples. These systematic biases raise an urgent need to identify their exact source, be it experimental [21], technical [17, 18] or computational [22]. Understanding the sources of biases is essential for the ongoing effort to mitigate and adjust for such biases from high-throughput collection and data compilation [23, 24].

Somatic mutations in cancer samples exhibit strong characteristics by the cancer type. An entirely opposite trend is expected for germline variants from healthy samples. There, the genomic characteristics signify the ethnic origin of the analyzed samples. By examining samples from the same population (i.e. Caucasians), we expect unified and cohesive genomic signals among samples and across cancer-types. Under such setting, it is easier to isolate the batch effect phenomenon, as we have shown here. However, we anticipate that the batch effect may also infiltrate, to some extent, into somatic mutation analyses, as suggested by our clinical analysis (Figure 5). However, due to the orders-of-magnitude higher variability in the number of somatic mutations observed among different cancer types, the batch effect is often masked, making it more challenging to identify. Many of the pan-cancer studies performed on TCGA data rely heavily on differences in the total number of somatic mutations among cancer types. Such studies might be skewed due to unaccounted sequencing batch effects. The identity of the genomic centers in which the blood samples were sequenced and the methodology used (i.e., the proportion between samples sequenced by HiSeq technology to data extracted from GeneArray) differ among cancer types and should be accounted for as well.

The reported TCGA batch effect has a broad range of implications. Our results demonstrate similarity among samples originating from the same sequencing center, compared to dissimilarity across samples from different sequencing centers. Our results reaffirm the encompassing nature of the sequencing batch effect that are not restricted to any particular cancer type from TCGA (Supplementary Figure S2).

In summary, the observed batch effects influence the number of variants per sample (Figure 1A), as well as the types of variants (Figure 1B), the number of variants at a per-gene resolution (Figure 2), and the distribution of variants within genes (Figure 3). They also drastically affect the majority of candidate genes annotated as predisposed for cancer (Figure 4) as well as other genes (Figure 6, Supplementary Table S2).

The batch effect described in this study is not restricted to the context of cancer, and may affect other human catalogues of WES germline variants. In the context of cancer, the pan-cancer studies are especially prone to batch effect that may lead to false discoveries and misinterpretation. Protocols for determining the identity and prevalence of somatic mutations from patient’s biopsy rely on having an accurate list of its germline variants. Developing methodologies to better control the inherent quantitative imbalances caused by batch effects is urgently and critically needed. Our results suggest that without batch effects correction, pan-cancer analysis cannot guarantee the precision required for personalized medicine. Current filters designed to remove batch effects from whole genome sequencing seem to impede the ability to detect true associations, and find new disease-associated variants [23,24]. In conclusion, the reported biases underlie the severe discrepancies in germline variants detection and analysis. Additionally, data from the different genomic centers may tamper with detection of somatic mutations, and therefore must be taken into consideration in any data driven pan-cancer analysis and interpretation.

## Materials and Methods

### Data resource

Approval for access BAM files and clinical data of TCGA cases was obtained from the database of Genotypes and Phenotypes (dbGaP) [25]. We selected a total of 2,241 blood derived healthy DNA samples with whole exome sequencing data (Supplementary Table S1). We limited the analysis to samples sequenced by the HiSeq-2000 Illumina technology. Aligned sequence data for normal samples in BAM file format and the accompanying metadata was downloaded from GDC portal [26].

### Germline variant calling

Variant calling was limited to exome regions only, as provided by UCSC GRCh38 reference genome [26]. We ran four different variant calling pipelines on each BAM file: GATK ‘HaplotypeCaller’ pipeline v3.5 [27], Atlas2 v1.4.3 [28], Freebayes [29] and Platypus v0.8.1 [30]. We filtered the results by their quality score. Samples with four complete VCF files (for each of the four pipelines) were unified; samples with missing or incomplete VCF files were discarded. Running this pipeline on a single BAM file took approximately 22 hours and produced a ^~^200MB unified VCF file.

### Comparing within-gene variant distributions

Many of the presented analyses required comparing the distributions of within-gene variant locations between cancer types (Figures 3,4,6). Within each gene, we collected all the called variants (from all samples), and partitioned them into six groups according to the cancer types they had originated from. We considered only the per-gene exomic locations of the variants (e.g. coordinates 0 to 8,300 for BRCA1, Figure 3). Denote by L_g,t_=(L_g,t_(1),..,L_g,t_(k_g,t_)) the collection of the gene exomic locations of all k_g,t_ called variants in a given gene g originating from samples of a given cancer type t (for example, if singleton germline variants were called at nucleotide positions 17, 65, and an additional variant was called at two individuals at position 183 of the KRAS transcript in SKCM samples, then L_KRAS,SKCM_=(17,65,183,183)). Note that the same locations, or even same variants, may appear multiple times in such a collection (e.g. if a variant is called in multiple samples). The empirical distributions of these variant location collections L_g,t_ are displayed in Figure 3 for all six cancer types within four selected genes.

In order to compare two cancer types t,s for a given gene g and obtain a p-value for the difference in the distributions of variants within that gene between the two cancer types, we applied a two-sided Kolmogorov-Smirnov (KS) test between the two (cumulative) empirical distributions of the collections, denoting the resulting p-values as p_g,(t,s)_=KS(L_g,t_,L_g,s_). These p-values are shown in supplementary table S3.

In order to obtain a final summary measure for the possible presence of batch effect within a gene (with respect to the distribution of variants along it), we took the ratio between the KS p-value of an intra sequencing center pair to the KS p-value of an inter sequencing center pair. Specifically, we defined the ratio r_g_=p_g,min_/p_g,max_ between the minimum of the p-values of BI-BI pairs p_g,min_=min (p_g,(SKCM,STAD)_, p_g,(SKCM,TCHA)_ p_g,(STAD,TCHA)_) to the maximum of the p-values of BI-WUGCS pairs p_g,max_=max(p_g,(SKCM,BRCA)_, p_g,(SKCM,UCEC)_, p_g,(STAD,BRCA)_, p_g,(STAD,UCEC)_, p_g,(TCHA,BRCA),_ p_g,(THCA,UCEC),_). We declared a gene to be possibly affected by the batch effect if r_g_>1. By taking a minimum-to-maximum ratio, we adopted a conservative criterion for the presence of the batch effect, requiring that all *between*-center p-values are smaller than all *within-centers* p-values. As reported, only 33% of the analyzed genes resulted a ratio r_g_< 1, indicating no batch effect.

## Supplementary Information

**Figure S1.**
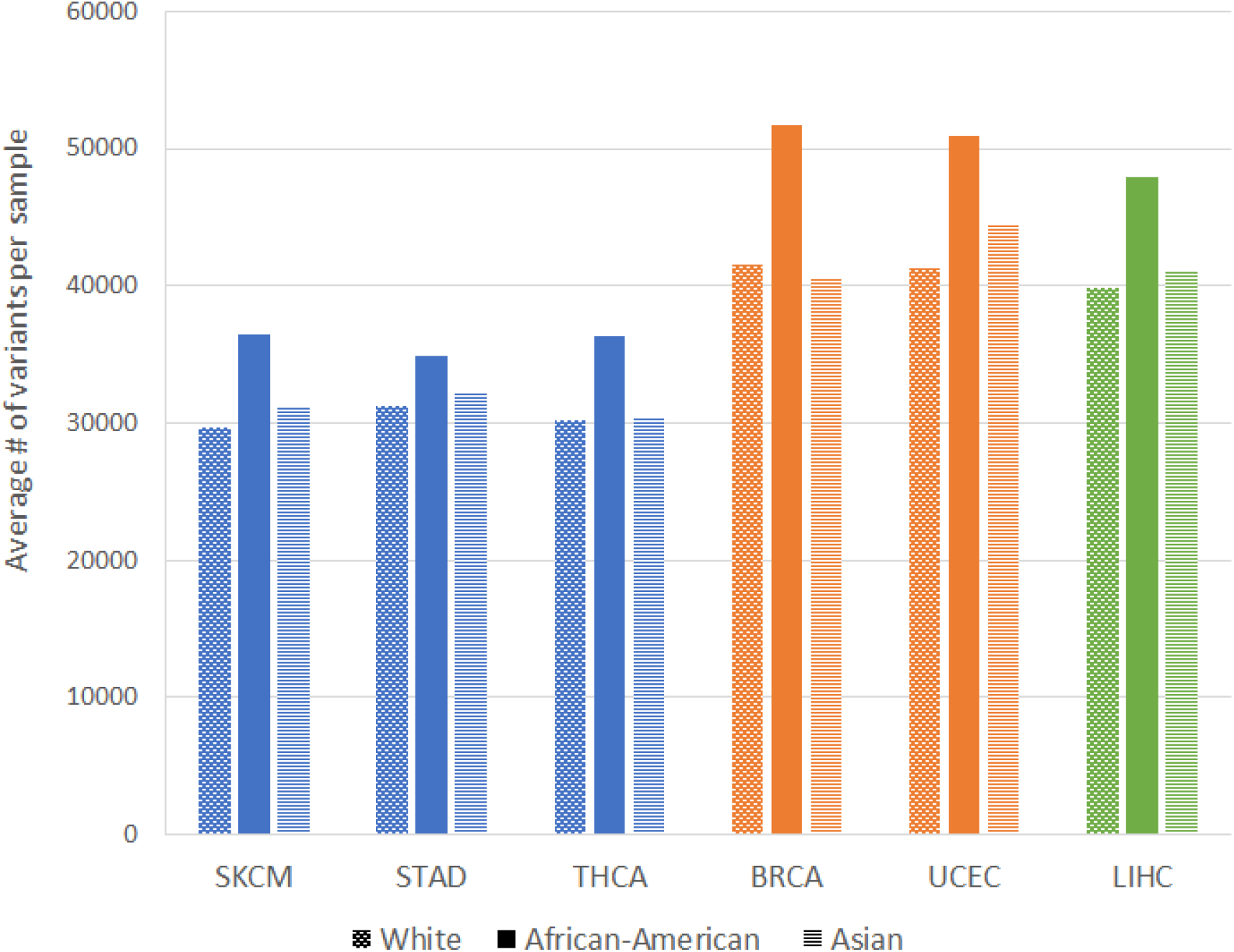
Number of exome variants per sample across ethnic groups and cancer types (data source: Supplementary Table S1)

**Figure S2.**
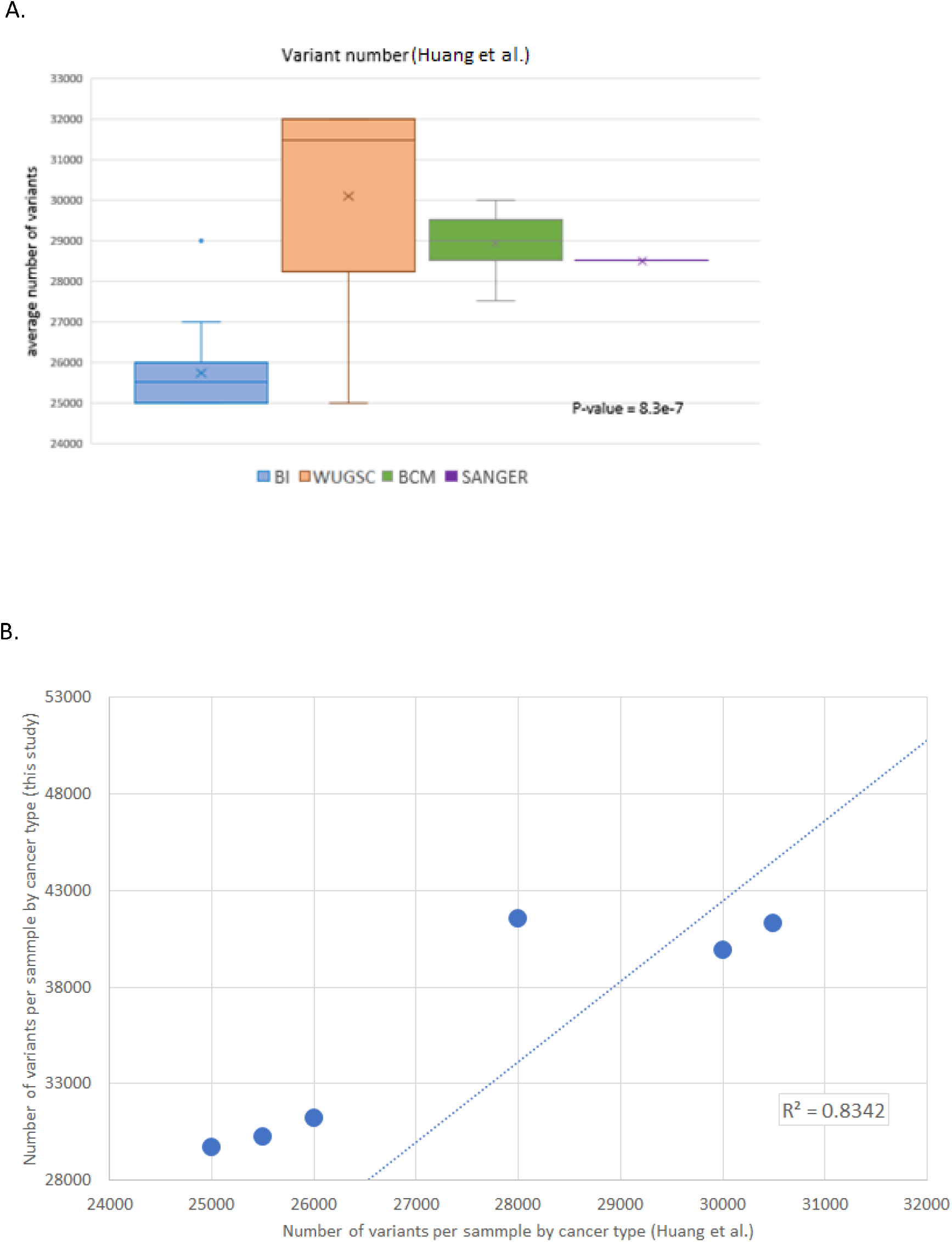
Average number of variants per sample based on an alternative variant calling pipeline (for all 33 cancer types) (A) Box plots show an average number of variants per sample across all 33 cancer types partitioned by the identity of the four sequencing centers. The data was extracted from another study (Huang et al. [7]; Supplementary Table S2). Boxplots are color-coded by the four sequencing centers that provided the analyzed data: Broad Institute (BI, blue), Washington University Genome Sequencing Center (WUGSC, orange) Baylor College of Medicine (BCM, green) and the Sanger Institute (Sanger, purple). A total of 16 cancer types were sequenced by BI, 9 by WUGSC, 7 by BCM, and 1 cancer type by the Sanger Institute. **(B)** Comparing the values of the 6 cancer types shared by the analysis in this study (y-axis, Supplementary Table S1) to the values in (Huang et al. [7]) (x-axis).

**Figure S3.**
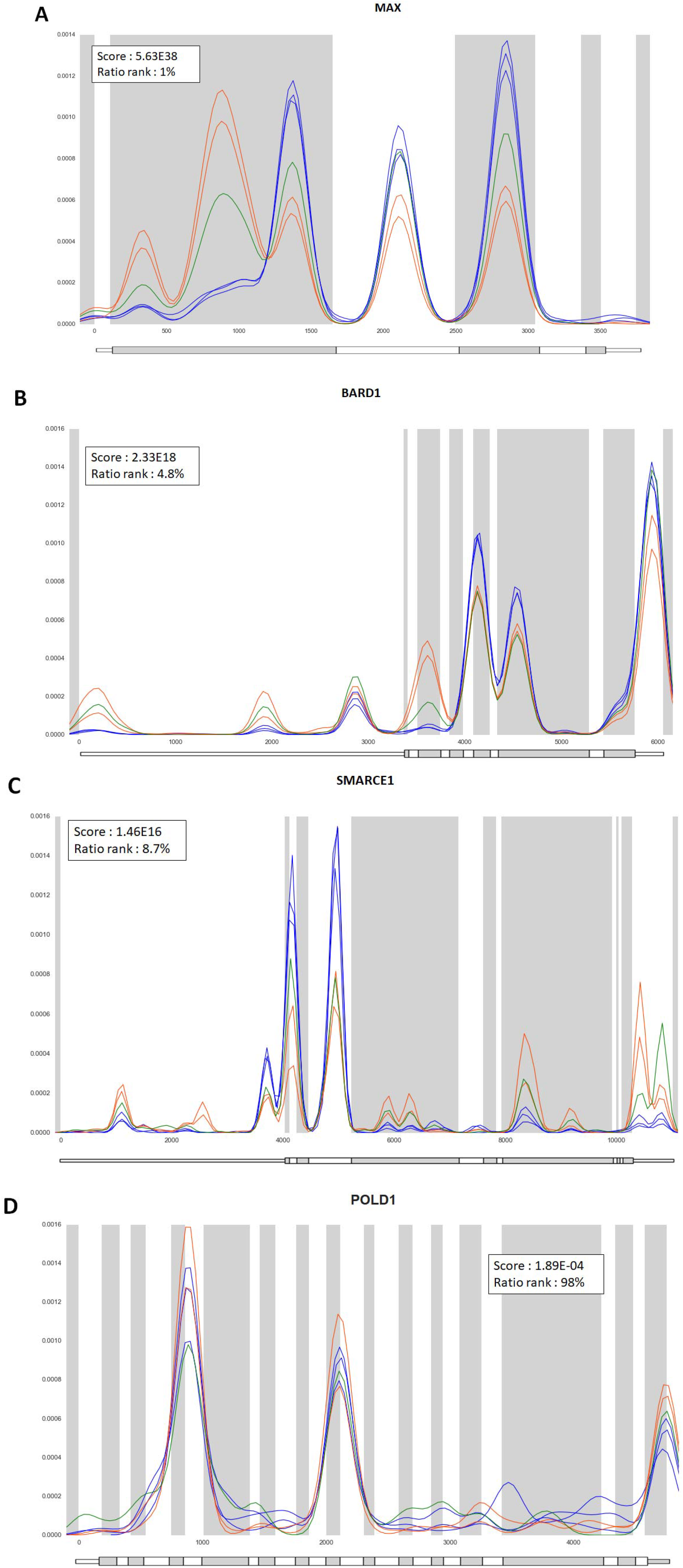
Gene exomic location distributions of germline variants within selected cancer. Empirical PDF of germline variant coordinates for four representative genes (see Figure 3 in main text). The genes shown are (A) MAX, (B) SMARCE1, (C) BARD1 and (D) POLD1. Each colored line represents the distribution for one of the 6 groups. SKCM, STAD, THCA are colored blue; BRCA, UCEC are in orange; LIHC is in green. The distributions per sequencing center are indicated by the score of the KS paired statistics (see Methods). The sensitivity rank is indicated by the percentage (lower percentage indicating higher sensitivity to the batch effect). KS p-values comparing variant distribution among all pairs per gene can

**Table S1.** Number of variants in exomes per sample across ethnic groups and cancer types

| **Cancer Type** | **White (Ave)** | **White (std)** | **African-American (Ave)** | **African-American (std)** | **Asian (Ave)** | **Asian (std)** |
| --- | --- | --- | --- | --- | --- | --- |
| **BRCA** | 41561.7 | 4770.1 | 51717.9 | 3304.6 | 40446.3 | 3825.4 |
| **LIHC** | 39884.5 | 3414 | 47976.3 | 5913.1 | 40996.4 | 2593 |
| **SKCM** | 29679.7 | 2382.5 | 36425 | - | 31134.2 | 4087.7 |
| **STAD** | 31228.9 | 3154.9 | 34824.2 | 1142.3 | 32259 | 3540.4 |
| **THCA** | 30251 | 1417.9 | 36294.3 | 1645.9 | 30368.6 | 1310.5 |
| **UCEC** | 41306.7 | 3825.2 | 50976.3 | 4114 | 44563.7 | 6770 |

**Table S2.**
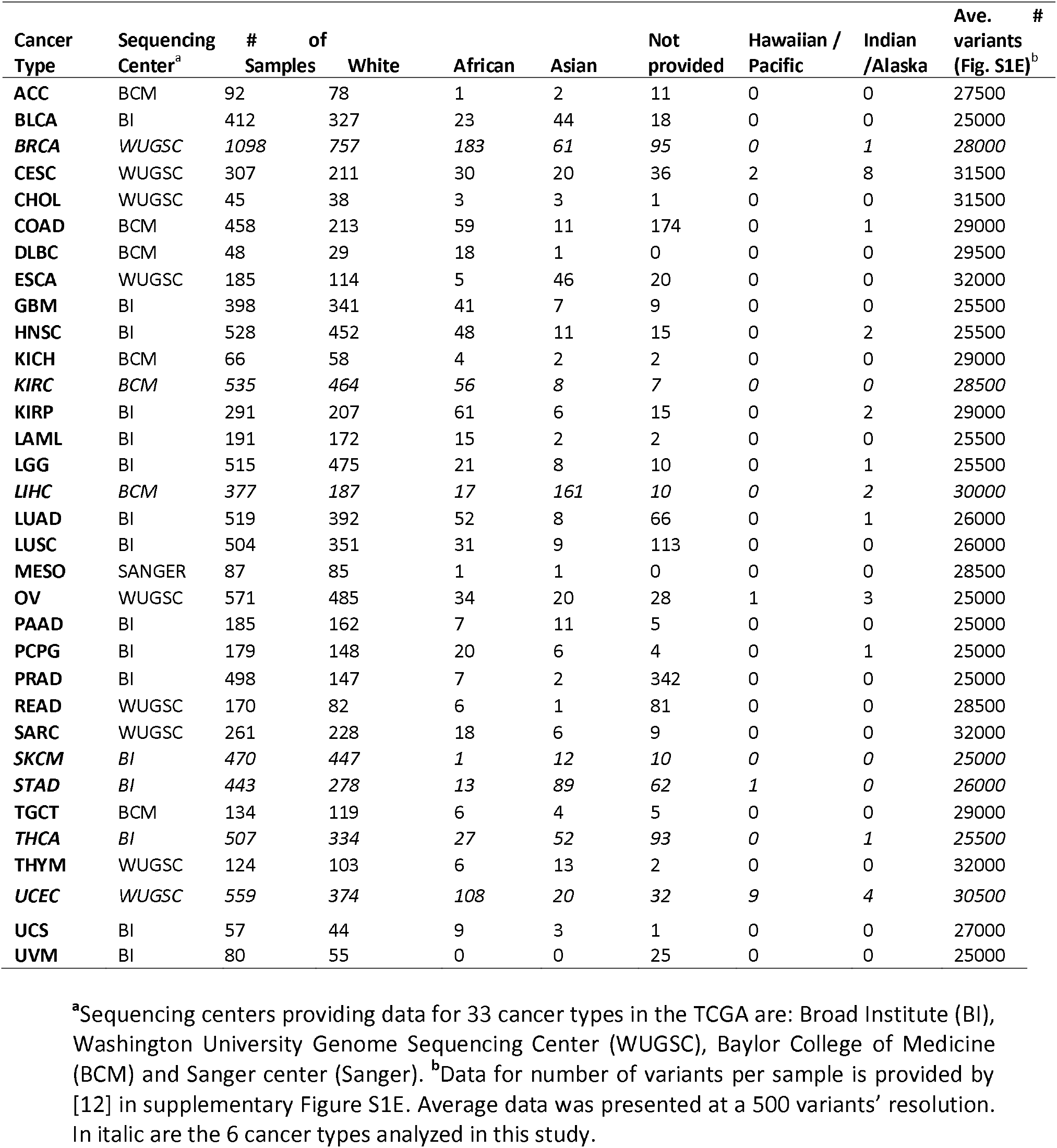
Cancer-type statistics derived from TCGA

**Table S3 -** Kolmogorov-Smirnov P-value per gene across all pairs of 6 cancer types

A measure of the batch distinctive variant distribution pattern is shown for the 104 CPG annotated by COSMIC (named “104 CPG”) and the entire genes (named “all genes”). The table lists all genes with at least a single variant among the compared groups.

## Abbreviations

BCM: Baylor college of medicine
BI: Broad Institute
GSC: Genome sequencing center
LoF: Loss of function
MPS: Massively parallel sequencing
SNV: Single nucleotide variant
KS: Kolmogorov-Smirnov
TCGA: The cancer genome atlas
TiTv: Transition-transversion
WUGSC: Washington university genome sequencing center
WES: Whole exome sequencing

## References

1. Lu C, Xie M, Wendl MC, Wang J, McLellan MD, Leiserson MD, Huang KL, Wyczalkowski MA, Jayasinghe R, Banerjee T, et al.: Patterns and functional implications of rare germline variants across 12 cancer types. Nat Commun 2015, 6:10086.

2. Rahman N: Realizing the promise of cancer predisposition genes. Nature 2014, 505:302–308.

3. Easton DF, Deffenbaugh AM, Pruss D, Frye C, Wenstrup RJ, Allen-Brady K, Tavtigian SV, Monteiro AN, Iversen ES, Couch FJ, Goldgar DE: A systematic genetic assessment of 1,433 sequence variants of unknown clinical significance in the BRCA1 and BRCA2 breast cancer-predisposition genes. Am J Hum Genet 2007, 81:873–883.

4. Risch HA, McLaughlin JR, Cole DE, Rosen B, Bradley L, Kwan E, Jack E, Vesprini DJ, Kuperstein G, Abrahamson JL, et al.: Prevalence and penetrance of germline BRCA1 and BRCA2 mutations in a population series of 649 women with ovarian cancer. Am J Hum Genet 2001, 68:700–710.

5. Gabai-Kapara E, Lahad A, Kaufman B, Friedman E, Segev S, Renbaum P, Beeri R, Gal M, Grinshpun-Cohen J, Djemal K, et al.: Population-based screening for breast and ovarian cancer risk due to BRCA1 and BRCA2. Proc Natl Acad Sci U S A 2014, 111:14205–14210.

6. Grant RC, Selander I, Connor AA, Selvarajah S, Borgida A, Briollais L, Petersen GM, Lerner-Ellis J, Holter S, Gallinger S: Prevalence of germline mutations in cancer predisposition genes in patients with pancreatic cancer. Gastroenterology 2015, 148:556–564.

7. Huang KL, Mashl RJ, Wu Y, Ritter DI, Wang J, Oh C, Paczkowska M, Reynolds S, Wyczalkowski MA, Oak N, et al.: Pathogenic Germline Variants in 10,389 Adult Cancers. Cell 2018, 173:355–370 e314.

8. van der Post RS, Vogelaar IP, Carneiro F, Guilford P, Huntsman D, Hoogerbrugge N, Caldas C, Schreiber KE, Hardwick RH, Ausems MG, et al.: Hereditary diffuse gastric cancer: updated clinical guidelines with an emphasis on germline CDH1 mutation carriers. J Med Genet 2015, 52:361–374.

9. Pearlman R, Frankel WL, Swanson B, Zhao W, Yilmaz A, Miller K, Bacher J, Bigley C, Nelsen L, Goodfellow PJ, et al.: Prevalence and Spectrum of Germline Cancer Susceptibility Gene Mutations Among Patients With Early-Onset Colorectal Cancer. JAMA Oncol 2017, 3:464–471.

10. Chubb D, Broderick P, Dobbins SE, Frampton M, Kinnersley B, Penegar S, Price A, Ma YP, Sherborne AL, Palles C, et al.: Rare disruptive mutations and their contribution to the heritable risk of colorectal cancer. Nat Commun 2016, 7:11883.

11. Wei R, Yao Y, Yang W, Zheng CH, Zhao M, Xia J: dbCPG: A web resource for cancer predisposition genes. Oncotarget 2016, 7:37803–37811.

12. Park S, Supek F, Lehner B: Systematic discovery of germline cancer predisposition genes through the identification of somatic second hits. Nat Commun 2018, 9:2601.

13. Cheng DT, Prasad M, Chekaluk Y, Benayed R, Sadowska J, Zehir A, Syed A, Wang YE, Somar J, Li Y, et al.: Comprehensive detection of germline variants by MSK-IMPACT, a clinical diagnostic platform for solid tumor molecular oncology and concurrent cancer predisposition testing. BMC Med Genomics 2017, 10:33.

14. Tomczak K, Czerwinska P, Wiznerowicz M: The Cancer Genome Atlas (TCGA): an immeasurable source of knowledge. Contemp Oncol (Pozn) 2015, 19:A68–77.

15. Lauss M, Visne I, Kriegner A, Ringner M, Jonsson G, Hoglund M: Monitoring of technical variation in quantitative high-throughput datasets. Cancer Inform 2013, 12:193–201.

16. Choi JH, Hong SE, Woo HG: Pan-cancer analysis of systematic batch effects on somatic sequence variations. BMC Bioinformatics 2017, 18:211.

17. Koire A, Katsonis P, Lichtarge O: Repurposing Germline Exomes of the Cancer Genome Atlas Demands a Cautious Approach and Sample-Specific Variant Filtering. Pac Symp Biocomput 2016, 21:207–218.

18. Buckley AR, Standish KA, Bhutani K, Ideker T, Lasken RS, Carter H, Harismendy O, Schork NJ: Pan-cancer analysis reveals technical artifacts in TCGA germline variant calls. BMC Genomics 2017, 18:458.

19. Forbes SA, Beare D, Boutselakis H, Bamford S, Bindal N, Tate J, Cole CG, Ward S, Dawson E, Ponting L, et al.: COSMIC: somatic cancer genetics at high-resolution. Nucleic Acids Res 2017, 45:D777–D783.

20. Zehir A, Benayed R, Shah RH, Syed A, Middha S, Kim HR, Srinivasan P, Gao J, Chakravarty D, Devlin SM, et al.: Mutational landscape of metastatic cancer revealed from prospective clinical sequencing of 10,000 patients. Nat Med 2017, 23:703–713.

21. Leek JT, Scharpf RB, Bravo HC, Simcha D, Langmead B, Johnson WE, Geman D, Baggerly K, Irizarry RA: Tackling the widespread and critical impact of batch effects in high-throughput data. Nat Rev Genet 2010, 11:733–739.

22. Zhang Z, Li H, Jiang S, Li R, Li W, Chen H, Bo X: A survey and evaluation of Web-based tools/databases for variant analysis of TCGA data. Brief Bioinform 2018.

23. Tom JA, Reeder J, Forrest WF, Graham RR, Hunkapiller J, Behrens TW, Bhangale TR: Identifying and mitigating batch effects in whole genome sequencing data. BMC Bioinformatics 2017, 18:351.

23. Zhang Y, Jenkins DF, Manimaran S, Johnson WE: Alternative empirical Bayes models for adjusting for batch effects in genomic studies. BMC Bioinformatics 2018, 19:262.

24. Grossman RL, Heath AP, Ferretti V, Varmus HE, Lowy DR, Kibbe WA, Staudt LM: Toward a Shared Vision for Cancer Genomic Data. N Engl J Med 2016, 375:1109–1112.

25. Wong KM, Langlais K, Tobias GS, Fletcher-Hoppe C, Krasnewich D, Leeds HS, Rodriguez LL, Godynskiy G, Schneider VA, Ramos EM, Sherry ST: The dbGaP data browser: a new tool for browsing dbGaP controlled-access genomic data. Nucleic Acids Res 2017, 45:D819–D826.

26. Tyner C, Barber GP, Casper J, Clawson H, Diekhans M, Eisenhart C, Fischer CM, Gibson D, Gonzalez JN, Guruvadoo L, et al.: The UCSC Genome Browser database: 2017 update. Nucleic Acids Res 2017, 45:D626–D634.

27. DePristo MA, Banks E, Poplin R, Garimella KV, Maguire JR, Hartl C, Philippakis AA, del Angel G, Rivas MA, Hanna M, et al.: A framework for variation discovery and genotyping using next-generation DNA sequencing data. Nat Genet 2011, 43:491–498.

28. Evani US, Challis D, Yu J, Jackson AR, Paithankar S, Bainbridge MN, Jakkamsetti A, Pham P, Coarfa C, Milosavljevic A, Yu F: Atlas2 Cloud: a framework for personal genome analysis in the cloud. BMC Genomics 2012, 13 Suppl 6:S19.

29. Blankenberg D, Von Kuster G, Bouvier E, Baker D, Afgan E, Stoler N, Galaxy T, Taylor J, Nekrutenko A: Dissemination of scientific software with Galaxy ToolShed. Genome Biol 2014, 15:403.

30. Rimmer A, Phan H, Mathieson I, Iqbal Z, Twigg SRF, Consortium WGS, Wilkie AOM, McVean G, Lunter G: Integrating mapping-, assembly- and haplotype-based approaches for calling variants in clinical sequencing applications. Nat Genet 2014, 46:912–918.

